# Dopamine vesicles are specified by mechanisms overriding canonical synaptic vesicle size constraints

**DOI:** 10.64898/2026.02.27.708430

**Authors:** Kenshiro Fujise, Nasser Karmali, Jaya Mishra, Himari Kimura, Tomoki Tsuji, Atsushi Saito, Nisha Mohd Rafiq

## Abstract

Dopaminergic neurons contain distinct synaptic vesicle (SV) populations that segregate dopamine from glutamate, but the molecular basis of this segregation remains unclear. Using an established in vitro SV reconstitution system in fibroblasts, we show that synaptogyrin family proteins—synaptogyrin 1, synaptogyrin 2 and synaptogyrin 3—but not synaptophysin, selectively associate with VMAT2-positive dopamine vesicles in the presence of synapsin. Despite the known role of synaptophysin and synaptogyrin family proteins in generating small SVs (40–50 nm), VMAT2 vesicles retain a larger size (50–80 nm) when co-expressed with synaptogyrins, while remaining segregated from synaptophysin-containing vesicles. SV2 family proteins (SV2A, SV2B, and SV2C) localize to both VMAT2- and synaptophysin-positive vesicle populations, with SV2C preferentially enriched on VMAT2-positive dopamine vesicles. Consistent with the selective dystrophy of SV2C/VMAT2/DAT-positive dopaminergic terminals in parkinsonism-associated SJ1^RQ^KI mice, iPSC-derived dopaminergic neurons carrying the same mutation accumulate small and enlarged SV-like organelles within autophagosomes, indicating defective SV clearance. Together, our findings suggest that dopamine vesicles are regulated by distinct trafficking mechanisms that override canonical SV size control, providing insight into the selective vulnerability of dopaminergic terminals in Parkinson’s disease.

## INTRODUCTION

Synaptic vesicles (SVs) are identified by their very small size in nerve terminals(De Camilli et al., 2001). Insight into mechanisms underlying this defining feature has come from reconstitution studies showing that ectopic expression of synaptophysin, one of the most abundant integral membrane proteins of SVs, together with the matrix protein synapsin, is sufficient to drive the formation of tightly clustered small vesicles within liquid-like condensates in fibroblasts (e.g., COS7 cells)(Milovanovic et al., 2018; Park et al., 2021). These vesicles closely resemble SVs observed at presynaptic terminals in the brain, at least with respect to size and protein composition.

Within this system, vesicular transporters for glutamate and GABA—VGLUTs and VGAT, respectively—coassemble with synaptophysin when coexpressed with synaptophysin and synapsin into small clear SV–like structures(Park et al., 2023). In contrast, VMAT2-containing vesicles that store dopamine display distinct behavior. These vesicles form larger vesicular clusters in the presence of synapsin and fail to assemble with synaptophysin, despite being positive for numerous other SV markers. Consistent with these findings, large-volume serial electron microscopy combined with proximity labeling of ventral tegmental area dopaminergic neurons has revealed that these neurons contain relatively few small SVs and that many boutons exhibit heterogeneous vesicle populations, including a substantial proportion of larger vesicles (>40–50 nm)(Lapios et al., 2025; Wildenberg et al., 2021). Differences between dopamine- and glutamate-containing vesicles are further supported by studies in mouse striatum, which have demonstrated marked distinctions between vesicles carrying VMAT2 and those containing VGLUTs, in part due to differences in recycling pathways(Fujise et al., 2025a; Fujise et al., 2025b; Onoa et al., 2010; Silm et al., 2019; Zhang et al., 2015). For example, experiments using pH-sensitive probes fused to either VMAT2 or VGLUT2 in mice lacking adaptor protein 3 (AP-3) showed a selective reduction in the formation of VMAT2-positive vesicles, while VGLUT2-positive vesicles were largely unaffected(Silm et al., 2019). In agreement with these observations, recent proteomic analyses have demonstrated that VGLUT2- and VMAT2-positive vesicles differ in protein composition, with synaptophysin more strongly associated with VGLUT2-positive vesicles than with VMAT2-positive vesicles(Asmerian et al., 2025).

Together, these findings indicate that dopaminergic neurons contain distinct SV populations in which dopamine is segregated from glutamate, yet the molecular mechanisms underlying this segregation remain poorly understood. In particular, it is unclear how vesicle identity and size are specified in dopamine vesicles, given their limited association with synaptophysin and their deviation from the canonical size of small SVs(Fujise et al., 2025b). Members of the synaptophysin and synaptogyrin family have been shown to be determinants of SV biogenesis and size(Park et al., 2024), raising the possibility that dopamine vesicles preferentially recruit alternative family members and that such recruitment may permit vesicle formation while bypassing canonical size constraints. In parallel, synaptic vesicle protein 2 (SV2) family members are broadly distributed across SVs and implicated in vesicle trafficking and transmitter stabilization(Buckley and Kelly, 1985; Janz et al., 1998), yet whether individual SV2 isoforms contribute to the molecular specialization of dopamine vesicles remains unknown. Of particular interest is SV2C, which is selectively enriched in dopaminergic brain regions and has been linked to dopaminergic dysfunction and Parkinsonian phenotypes in both genetic and disease models(Dunn et al., 2017; Foo et al., 2020; Ng et al., 2023). These observations raise the question of whether selective association of SV2 family members, and SV2C in particular, contributes to the identity, trafficking, and vulnerability of dopamine vesicles.

In this study, we combine SV reconstitution with analyses of dopaminergic neurons to define how synaptophysin family proteins (synaptophysin, synaptogyrin 1, synaptogyrin 2, and synaptogyrin 3) and SV2 isoforms (SV2A, SV2B, and SV2C) selectively associate with VMAT2-positive vesicles and regulate their segregation, size, and organization. We further examine the size of SVs in iPSC-derived dopaminergic neurons carrying a Parkinson’s disease(PD)–associated SJ1 mutation, which exhibit defects in SV endocytosis and the accumulation of both small and enlarged SV-like vesicles.

## RESULTS

### Distinct localizations of synaptophysin and synaptogyrins on VMAT2-positive SVs

We utilized an in cellulo SV reconstitution system that has been shown to generate clusters of SV-like organelles in fibroblastic cells (COS7 cells) through the exogenous expression of synapsin and synaptophysin(Park et al., 2021). In this system, these vesicles self-organize into liquid-like condensates that appear as large droplets by fluorescence microscopy. Consistent with previous observations, co-expression of untagged synaptophysin and mCherry-synapsin in COS7 cells resulted in the formation of droplet-like condensates (Figure 1A). In contrast, combined expression of the dopamine transporter, VMAT2 (VMAT2-GFP), with synaptophysin and mCherry-synapsin led to the formation of condensates that segregated into distinct phases, with VMAT2-positive vesicles separating from synaptophysin-containing condensates (Figure 1B), as we have previously reported (Fujise et al., 2025b).

**Figure 1.**
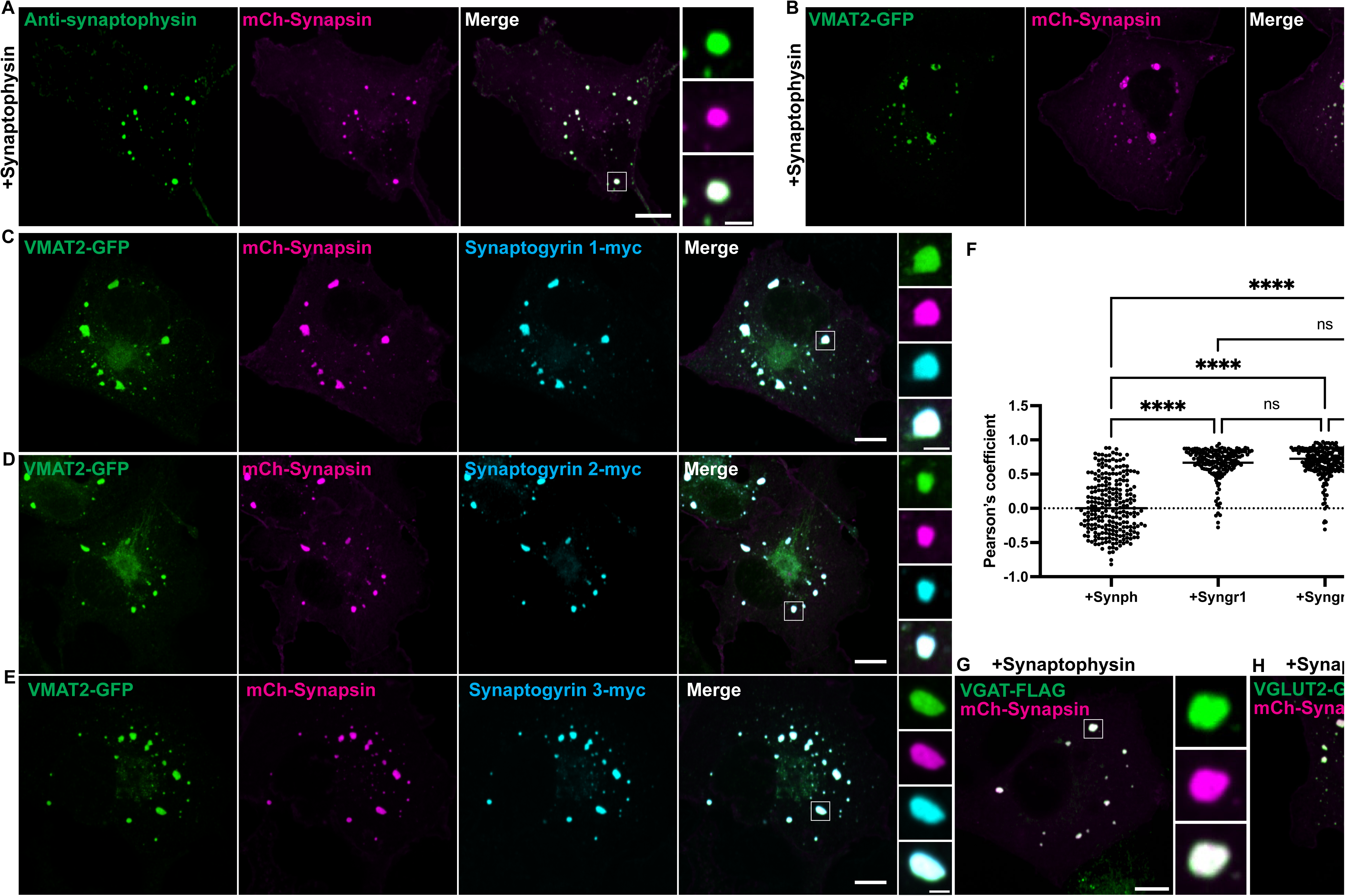
Synaptogyrin family proteins, but not synaptophysin, colocalize with VMAT2-positive SV-like organelles. (A-B) Representative fluorescence images of COS7 cells transfected with (A) untagged synaptophysin (green) and mCherry-synapsin (magenta) or (B) untagged synaptophysin, VMAT2-GFP (green) and mCherry-synapsin (magenta). Untagged synaptophysin was detected by immunofluorescence using an antibody against synaptophysin in A. (C-E) Representative fluorescence images of COS7 cells transfected with VMAT2-GFP (green) and mCherry-synapsin (magenta), together with either synaptogyrin 1–Myc (cyan; C), synaptogyrin 2–Myc (cyan; D), or synaptogyrin 3–Myc (cyan; E). All three synaptogyrin family proteins show co-localization with VMAT2–synapsin condensates. The Myc signal of synaptogyrin 1, synaptogyrin 2, and synaptogyrin 3 was detected by immunofluorescence using an anti-Myc antibody. (F) Quantification of colocalization between synaptophysin, synaptogyrin 1, synaptogyrin 2, or synaptogyrin 3 and VMAT2–synapsin condensates, expressed as Pearson’s correlation coefficient (n = 3 independent experiments). ns, not significant; ****, p < 0.0001 (one-way ANOVA). (G-H) Co-expression of the GABA transporter VGAT-FLAG (G) or the glutamate transporter VGLUT2-GFP (H) results in colocalization with untagged synaptophysin-positive vesicles, in contrast to VMAT2. Scale bars, 10 μm; inset 2.5 μm.

We next sought to determine whether other members of the synaptophysin family—synaptogyrin 1, synaptogyrin 2, and synaptogyrin 3—associate differently with VMAT2-containing vesicles and whether their recruitment influences vesicle organization within VMAT2–synapsin condensates. Each of these proteins, when expressed individually with synapsin, has been shown to be sufficient to generate clusters of small vesicles comparable in size to synaptophysin-positive vesicles(Park et al., 2024). In contrast to synaptophysin, however, all three synaptogyrins (Synatogyrin 1-Myc, Synatogyrin 2-Myc, Synatogyrin 3-Myc), co-assembled with vesicles within VMAT2–synapsin (VMAT2-GFP, mCherry-synapsin) condensates (Figure 1C–F). It has been previously proposed that the C-terminal regions of synaptophysin and synaptogyrin family proteins carry a net negative charge that interacts with the highly basic C-terminal tail of synapsin(Park et al., 2024), providing a plausible mechanism for SV clustering (Supplementary Figure 1A-C). Consistent with this model, analysis of VMAT2 revealed that its C-terminal region is also negatively charged (pI = 4.695; net charge at pH 7.4 = −4.935), suggesting a potential electrostatic interaction with the basic C-terminal domain of synapsin that may contribute to the formation of VMAT2-containing vesicle clusters.

In addition, consistent with our previous findings(Fujise et al., 2025b), VGAT (VGAT-FLAG) and VGLUT2 (VGLUT2-GFP)—but not VMAT2—co-assembled with synaptophysin–synapsin condensates when co-expressed in COS7 cells (Figure 1G,H). Together, these results indicate that, despite their close structural similarity to synaptophysin, synaptogyrin family members exhibit distinct behavior in the context of VMAT2-positive vesicles, supporting differential roles for synaptophysin and synaptogyrins in the biogenesis and clustering of dopamine-containing SVs.

### Size regulation of VMAT2-positive SVs by synaptogyrins

Next, we investigated whether synaptogyrins influence the size of dopamine-containing SVs. Synaptogyrin-induced condensates have previously been shown to consist of clusters of SV-like vesicles with an average diameter of ∼40 nm (Figure 2A; Park et al., 2024). Among synaptogyrin family members, synaptogyrin 3 is of particular interest, as it has been reported to facilitate dopamine uptake through interactions with DAT and VMAT2 (Egana et al., 2009) and is highly expressed in dopaminergic nerve terminals(Hobson et al., 2022). We therefore focused on synaptogyrin 3 to examine whether its recruitment alters the size of VMAT2-positive vesicles.

**Figure 2.**
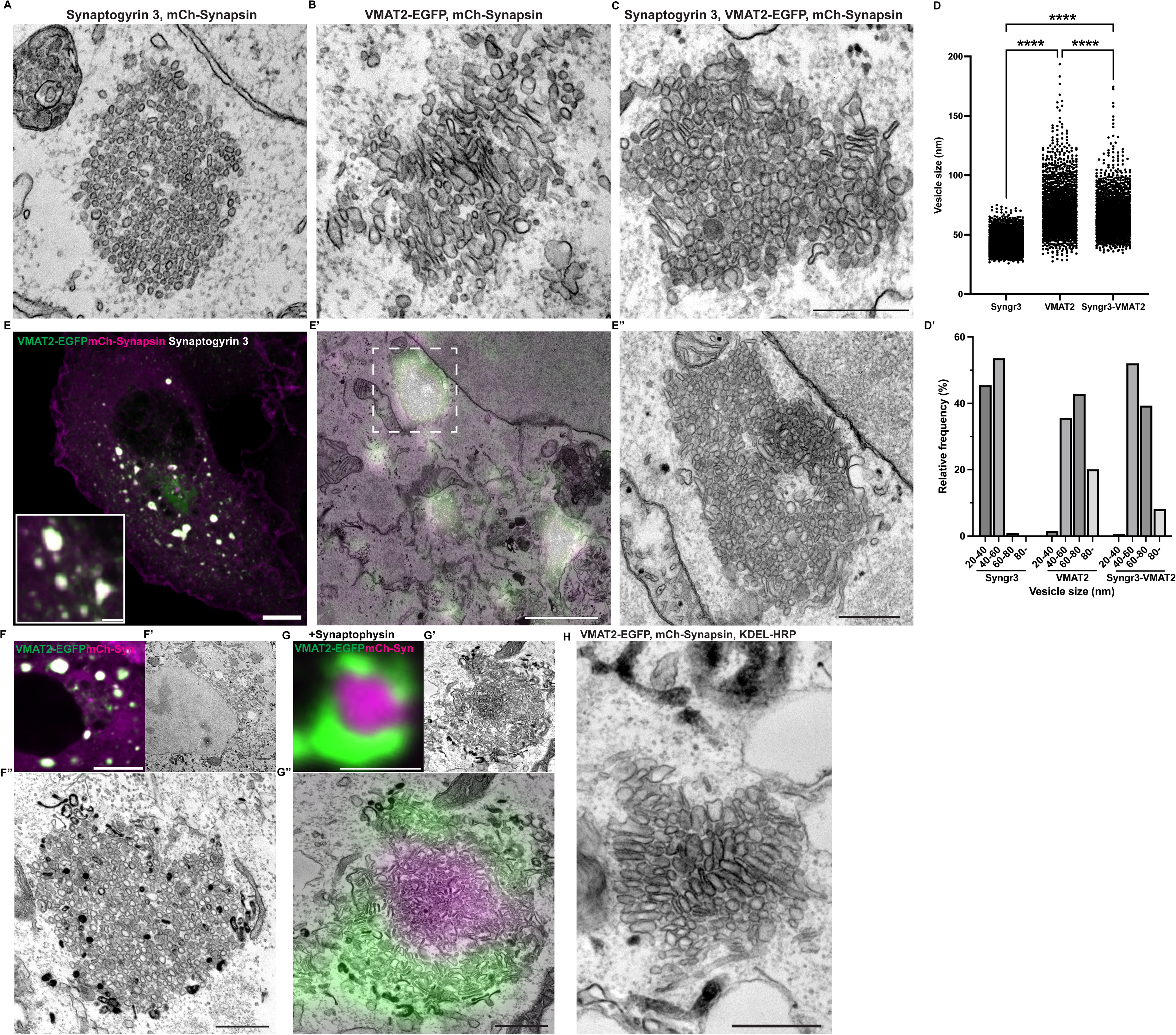
Synaptogyrin modulates VMAT2-positive vesicles without restoring canonical SV size. (A, B) Representative electron micrographs show clusters of vesicles with canonical SV size when untagged synaptogyrin 3 is co-expressed with mCherry-synapsin (A), in contrast to the large and pleiomorphic vesicles observed in VMAT2–synapsin condensates (B). (C) Co-expression of untagged synaptogyrin 3, mCherry-synapsin, and VMAT2-GFP resulted in vesicles that were larger than canonical SVs (40–50 nm) but more uniform in size and more circular in shape than vesicles formed by VMAT2-GFP and mCherry-synapsin alone (B). Scale bar, 500 nm. (D, D’) Quantification of individual vesicle diameters from vesicle clusters shown in (A–C), presented as a dot plot (D; ANOVA) and frequency histogram (D’). ns, not significant; ****, p < 0.0001 (one-way ANOVA). The dot plot is represented as mean ± SD pooled from ≥ 2351 vesicles in vesicle clusters of three different cells. ((E) Correlative light–electron microscopy (CLEM) of COS7 cells co-transfected with untagged synaptogyrin 3, VMAT2-GFP (green), and mCherry-synapsin (magenta). Scale bars, 10 μm; inset 2 μm. The fluorescence image of the inset is overlaid onto the corresponding transmission electron micrograph (TEM) image (E’). Scale bar, 2 μm. (E’’) Higher-magnification view of a representative cluster containing enlarged vesicles. Scale bar, 500 nm. (F-G) CLEM images of cropped regions from COS7 cells co-transfected with VMAT2-GFP and mCherry-synapsin (F) or with VMAT2-GFP, untagged synaptophysin, and mCherry-synapsin (G), following incubation with CTX-HRP for 1 h. The corresponding electron micrographs are shown in (F’) and (G’). Dark electron-dense staining indicates HRP-reactive labeling of vesicles originating from the plasma membrane in both VMAT2-only vesicles (F’’) and vesicles containing both synaptophysin and VMAT2 in distinct phases (G’’). Synaptophysin-positive vesicles are readily identified by their small size and strong mCherry-synapsin signal, whereas VMAT2-positive vesicles exhibit weaker synapsin fluorescence and larger size. Scale bars for fluorescence image: 10 μm; EM (low magnification): 2 μm; EM(high magnification): 500 nm. (H) Representative electron micrograph of COS7 cells co-transfected with VMAT2-GFP and mCherry-synapsin following incubation with ER-targeted KDEL–HRP for 2 h. VMAT2–synapsin clusters lack electron-dense HRP staining,

To test this hypothesis, synaptogyrin 3 was co-expressed with mCherry-synapsin in COS7 cells, either in the presence or absence of VMAT2-GFP, and vesicle size within the resulting condensates was analyzed by electron microscopy. Synaptogyrin 3–synapsin condensates were indistinguishable from synaptophysin–synapsin condensates, forming clusters of small vesicles in the canonical SV size range (41.6 ± 6.7 nm; Figure 2A), consistent with previous report (Park et al., 2024). As we have previously demonstrated, larger vesicles are observed when VMAT2 is co-expressed with synapsin, as also demonstrated here (68.6 ± 19.13 nm; Figure 2B). In contrast, co-expression of VMAT2-GFP with synaptogyrin 3 and mCherry-synapsin resulted in the formation of larger vesicles than those formed by synaptogyrin 3-synapsin, but modestly smaller than those in the VMAT2–synapsin condition, with an average diameter of 62.1 ± 12.8 nm (Figure 2C-D’). To confirm that these condensates corresponded to clusters of enlarged vesicles, we performed correlative light and electron microscopy (CLEM). CLEM analysis revealed that VMAT2–synapsin–synaptogyrin 3 condensates consistently comprised clusters of large vesicles across all condensates observed in COS7 cells (Figure 2E-E’’). Together, these results indicate that synaptogyrin family proteins can modulate vesicle size but are not sufficient to impose canonical SV size constraints on VMAT2-positive vesicles. Instead, the trafficking pathway associated with VMAT2 appears to dominate vesicle sizing, with synaptogyrins exerting a secondary modulatory effect.

Given the irregular size and morphology of VMAT2-positive vesicles and prior reports suggesting that VMAT2-containing membranes may derive from the endoplasmic reticulum(Nirenberg et al., 1996; Nirenberg et al., 1995), we examined whether these vesicles originate from distinct membrane trafficking pathways. We assessed their endocytic origin using cholera toxin–horseradish peroxidase (CTX-HRP) uptake in combination with CLEM. Both VMAT2-positive vesicles formed in the presence of synapsin alone and those formed upon co-expression with synaptophysin were labeled by CTX-HRP (Figure 2F–G’’), indicating that these vesicle populations have endocytic origins. To further evaluate a potential contribution from the endoplasmic reticulum (ER), cells were loaded with KDEL-HRP to label the ER lumen. While ER membranes were electron-dense under these conditions, VMAT2-GFP–positive vesicles lacked HRP labeling in the presence of synapsin, and were readily distinguishable from ER structures by electron microscopy(Figure 2H). Together, these observations support an endocytic origin for VMAT2-associated vesicles formed under these conditions.

### Differential localization of SV2 isoforms to VMAT2- and synaptophysin-positive vesicles

SV2 proteins are large glycosylated transmembrane vesicle proteins with 12 predicted transmembrane domains and extensive luminal loops, a conserved architecture thought to support interactions with vesicle membranes and trafficking machinery(Janz et al., 1998). SV2 proteins exhibit distinct expression patterns across brain regions: SV2A is ubiquitously expressed, SV2B shows regionally restricted expression (e.g., cortex and thalamus), and SV2C is predominantly expressed in the dopaminergic-striatal regions(Dunn et al., 2017). Consistent with this distribution, multiple proteomic analyses of dopaminergic nerve terminals and striatal synaptosomes have repeatedly demonstrated enrichment of SV2C in SVs associated with dopamine transmission (Hobson et al., 2022; Oostrum et al., 2023; Paget-Blanc et al., 2022). To investigate whether these differences in expression reflect intrinsic vesicle-sorting preferences, we examined the localization of SV2 family members—SV2A, SV2B, and SV2C—which share high structural homology, using the SV reconstitution assay in COS7 cells.

To assess whether SV2 isoforms are sufficient to drive synapsin-dependent vesicle clustering, we expressed SV2A, SV2B, or SV2C in COS7 cells, either alone or together with synapsin. All three SV2 isoforms localized to Golgi-associated and vesicular puncta when expressed alone(Supplementary Figure 2A-C), but none induced the formation of synapsin-dependent condensates when co-expressed with mCherry-synapsin (Supplementary Figure 2D-F). We then assessed SV2 recruitment to synaptophysin–synapsin and VMAT2–synapsin condensates individually. All three SV2 isoforms were efficiently recruited to condensates induced by synaptophysin and synapsin and showed extensive co-localization with these structures (Figure 3A–C). Similarly, VMAT2–synapsin condensates recruited SV2A, SV2B, and SV2C, with no obvious differences in localization when assessed individually, as judged by fluorescence overlap in high-magnification images (Figure 3D–F).

**Figure 3.**
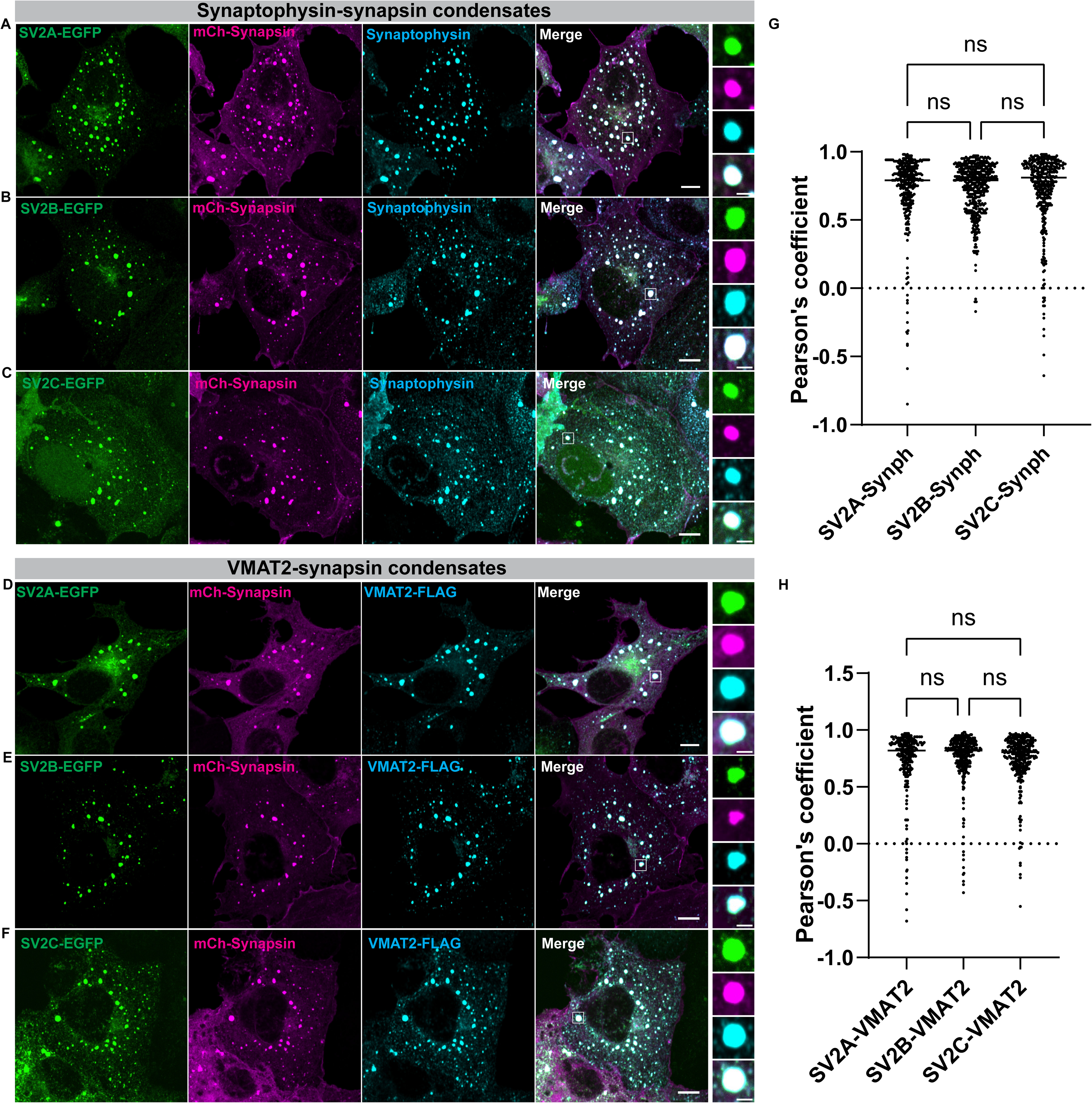
SV2 family proteins co-localize with vesicles in both synaptophysin–synapsin and VMAT2–synapsin condensates. Representative fluorescence images of COS7 cells transfected with (A–C) untagged synaptophysin (cyan) and mCherry-synapsin (magenta), or (D–F) VMAT2-FLAG (cyan) and mCherry-synapsin (magenta), together with either SV2A-GFP (green; A,D), SV2B-GFP (green; B,E), or SV2C-GFP (green; C,F). Note the strong white signal in the cropped condensates, indicating overlap among all channels. The FLAG signal of VMAT2-FLAG was detected by immunofluorescence using an anti-FLAG antibody. Scale bars, 10 µm; inset, 2.5 µm. (G, H) Quantification of the colocalization of SV2A, SV2B, and SV2C with synaptophysin (G) or VMAT2 (H) within synaptophysin–synapsin (G) and VMAT2–synapsin (H) condensates, respectively, expressed as Pearson’s correlation coefficients (n = 3 independent experiments). ns, not significant.

We therefore next asked whether SV2 isoforms exhibit preferential localization when both vesicle populations are present simultaneously (Figure 4). To this end, we examined SV2 distribution within segregated condensates formed by co-expression of VMAT2-FLAG, untagged synaptophysin, and mCherry-synapsin. In this configuration, synaptophysin–synapsin condensates were identified by increased synapsin fluorescence in the absence of VMAT2-FLAG signal. Under these conditions, SV2A did not show a preference for synaptophysin- or VMAT2-positive vesicles and was recruited to both types of condensates. In contrast to SV2B, which displayed a relative preference for synaptophysin-positive condensates (Figure 4A–D and G), SV2C showed a pronounced enrichment in VMAT2-positive condensates when given the choice between the two vesicle populations (Figure 4E-G). This selective enrichment of SV2C in VMAT2-positive condensates is consistent with its preferential representation in dopaminergic vesicle proteomes and supports the view that the in vitro reconstitution system captures key aspects of SV organization in dopaminergic neurons.

**Figure 4.**
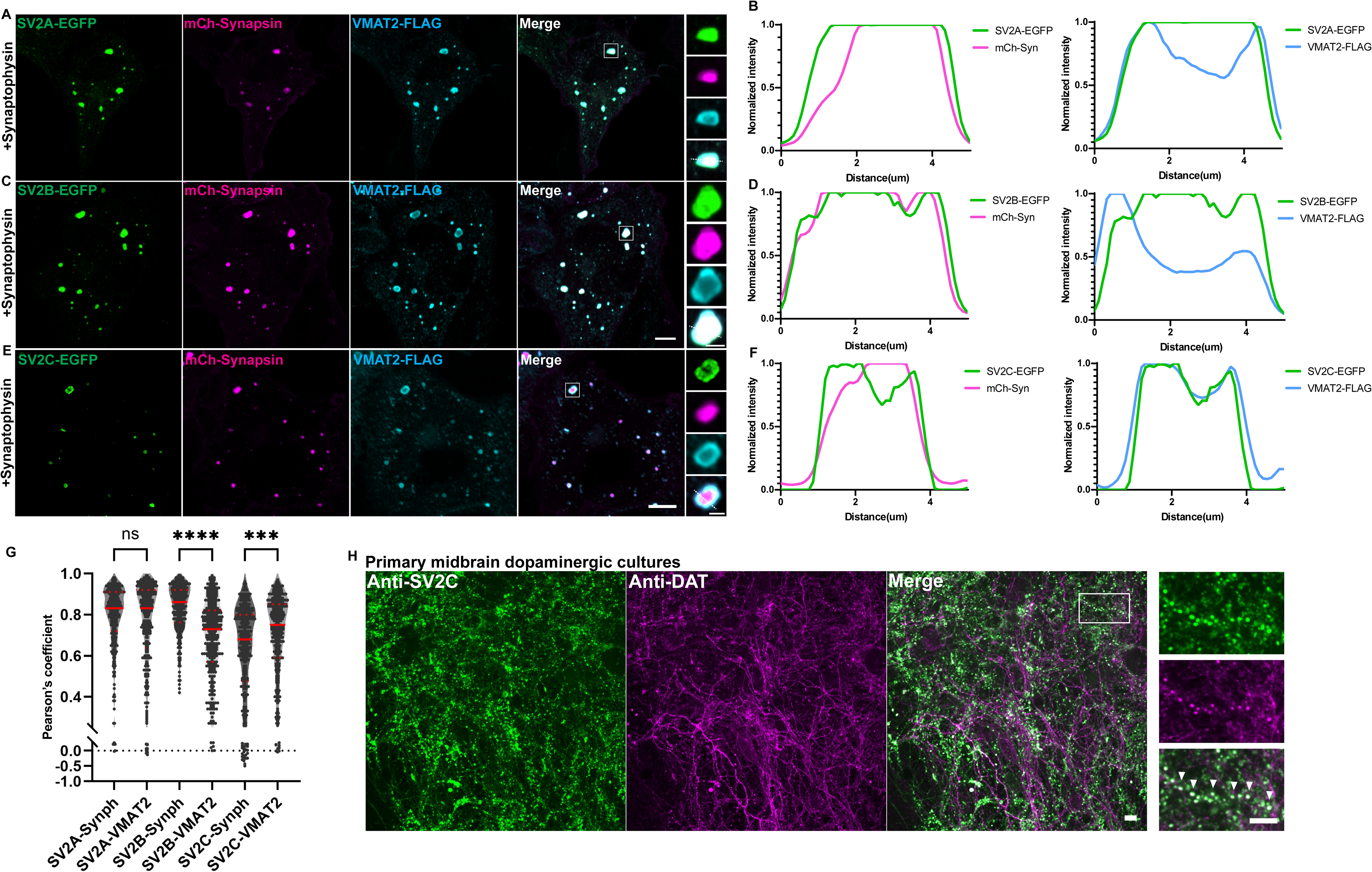
SV2C preferentially associates with VMAT2–synapsin condensates when co-expressed with synaptophysin. Representative fluorescence images of COS7 cells transfected with untagged synaptophysin and mCherry-synapsin (magenta), together with VMAT2-FLAG (cyan) and either SV2A-GFP (green; A), SV2B-GFP (green; C), or SV2C-GFP (green; E). Corresponding line-scan intensity profiles from the cropped condensates are shown in (B) for SV2A, (D) for SV2B, and (F) for SV2C. Synaptophysin–synapsin condensates were identified by high synapsin fluorescence (left graph panels), whereas VMAT2 condensates were identified by VMAT2-FLAG signal (right graph panels) and lower synapsin intensity. Note that SV2A localizes to both synaptophysin- and VMAT2-positive condensates, whereas SV2C shows a clear preference for VMAT2-positive condensates (F, right graph panel). The FLAG signal of VMAT2-FLAG was detected by immunofluorescence using an anti-FLAG antibody. (G) Quantification of the colocalization of SV2A, SV2B, and SV2C with synaptophysin or VMAT2 within mixed synaptophysin–synapsin and VMAT2–synapsin condensates, expressed as Pearson’s correlation coefficients (n = 3 independent experiments). ns, not significant; ***, p=0.0001, ****, p < 0.0001 (Kruskal-Wallis test). (H) Immunofluorescence images of midbrain dopaminergic neuron cultures at 20 days in vitro show numerous SV2C-positive puncta (green) overlapping with DAT-positive presynaptic terminals (magenta). SV2C and DAT were detected by immunofluorescence using antibodies against them. Scale bars, 20 µm; inset, 5 µm.

To gain insight into the molecular basis of this selectivity, we examined the predicted structures of SV2 isoforms using AlphaFold. All three SV2 proteins display highly similar overall topology (∼80% similarity), consisting of 12 transmembrane helices, with the cytosolic N-terminal regions showing the greatest sequence and structural variation (Supplementary Figure 3A–D). Notably, SV2A and SV2C share greater similarity within their N-terminal regions compared with SV2B (Supplementary Figure D). Whereas the N-termini of SV2A and SV2C comprise two helices connected by a disordered segment, the N-terminus of SV2B is largely disordered and contains only a single predicted helix (Supplementary Figure 3A’–C’). Whether these differences in the cytosolic N-terminal regions contribute to the reduced association of SV2B with VMAT2-positive vesicles relative to SV2C remains unclear, and future protein-based chimera studies will be required to address this possibility.

Consistent with these observations in the reconstitution system, immunofluorescence analysis of primary dopaminergic neurons revealed co-localization of SV2C with the dopamine transporter DAT, a marker of dopaminergic presynaptic terminals (Figure 4H). This finding further supports the association of SV2C with dopaminergic vesicles, in agreement with previous proteomic studies(Asmerian et al., 2025; Hobson et al., 2022; Oostrum et al., 2023; Paget-Blanc et al., 2022). Together, these results indicate that SV2 isoforms exhibit differential vesicle-sorting preferences, with SV2C selectively associating with VMAT2-positive vesicle populations. Moreover, vesicle clusters induced by VMAT2 and synaptophysin possess distinct molecular compositions, supporting the notion that these vesicle populations are organized through independent trafficking pathways(Asmerian et al., 2025).

### PD-associated SJ1 mutation disrupt dopamine vesicle homeostasis and clearance

To assess whether synaptic dysfunction is a key feature of PD, we examined large-scale transcriptomic datasets from human PD caudate and putamen (dorsal striatum) samples (Irmady et al., 2023). Differential expression analyses revealed broad downregulation of genes associated with synaptic vesicle function and dopaminergic terminals, including synaptogyrin family proteins, SV2C, VMAT2, and endocytic recycling factors linked to familial PD such as synaptojanin-1 (SJ1) and DNAJC6/auxilin (Figure 5A). Gene Ontology (GO) analyses further identified significant enrichment of pathways related to synaptic vesicle trafficking, dopaminergic synapses, endocytosis, and autophagy among the downregulated genes (Figure 5B). Consistent with these findings, SynGO analysis (Koopmans et al., 2019) demonstrated that genes associated with the presynaptic membrane and synaptic-related processes were preferentially reduced in PD brains (Figure 5C). Together, these observations suggest that defects in synaptic vesicle organization and recycling are prominent features of PD dopaminergic terminals.

**Figure 5.**
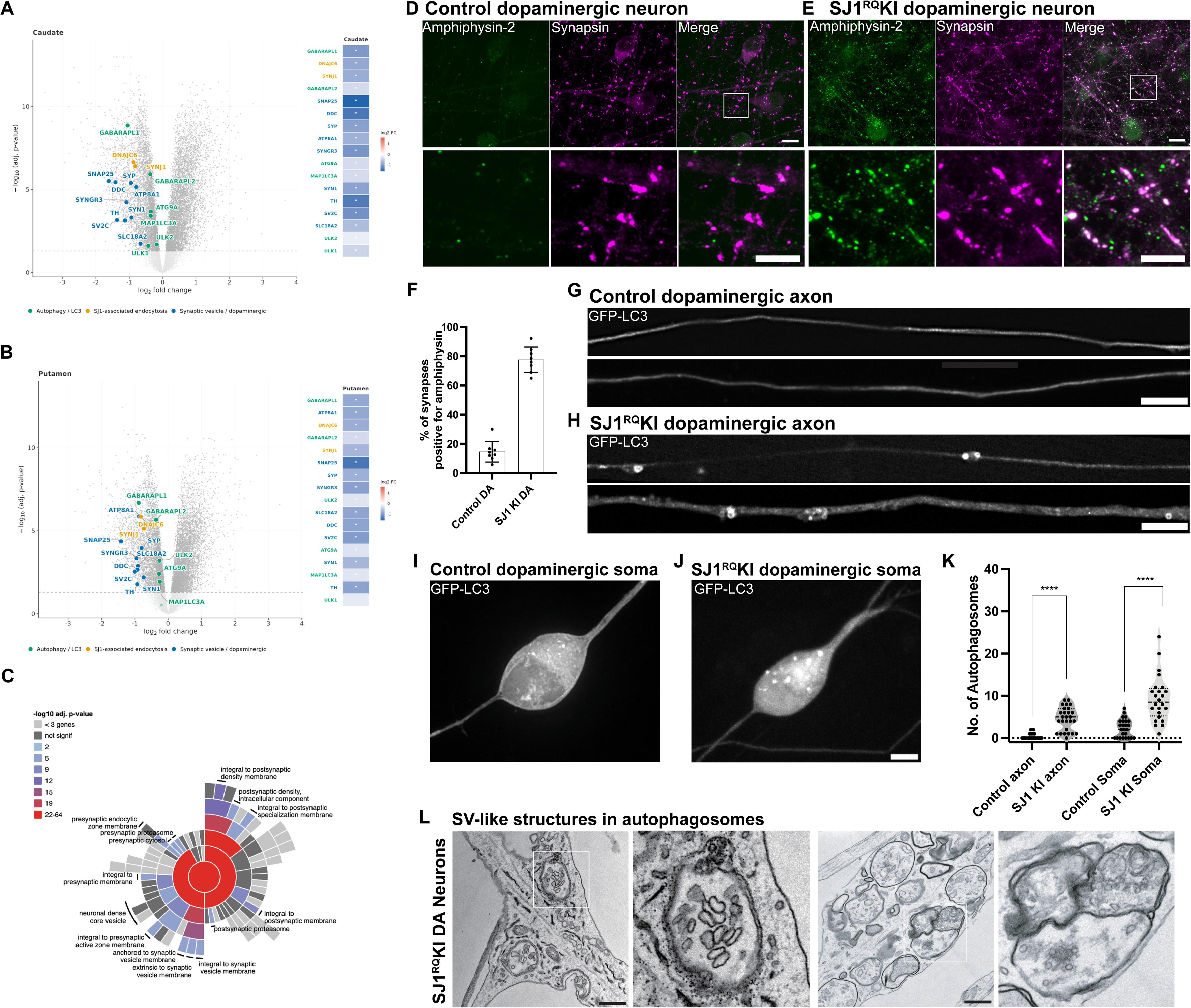
Small and large SV-like structures accumulate within autophagosomes in SJ1^RQ^ knock-in dopaminergic neurons. (A-C) Transcriptome analysis of postmortem striatum from Parkinson’s disease patients and neurologically normal controls (Irmady et al., 2023). (A) Volcano plots displaying differential gene expression in the caudate (A) and putamen (B). Genes are color-coded by functional category: synaptic vesicle / dopaminergic (blue), SJ1-associated endocytosis (orange), and autophagy / LC3 (green). Filled circles indicate significantly differentially expressed genes (adj. p-value < 0.05). Inset heatmaps show negative log2 fold changes for all genes of interest across both regions, ordered by adjusted p-value; asterisks indicate significance (adj. p-value < 0.05). (C) SynGo sunburst plot identifying downregulated genes are enriched in the presynaptic membrane and synaptic processes (red sectors). (D–F) SJ1^RQ^KI iPSC-derived dopaminergic neurons show presynaptic clustering of amphiphysin-2. (D, E) Representative fluorescence images of control (D) and SJ1^RQ^KI (E) dopaminergic neurons (days 55–65 in culture) immunolabeled with antibodies against amphiphysin-2 (green) and synapsin, a presynaptic marker (magenta). Scale bars, 10 µm. Higher-magnification views of the boxed regions are shown below each panel. Scale bars, 5 µm. Note the marked increase in amphiphysin-2 immunoreactivity overlapping with synapsin-positive structures in SJ1^RQ^KI dopaminergic neurons relative to controls. (F) Quantification of the percentage of synapses showing amphiphysin-2 clustering in the conditions shown in (D, E), presented as mean ± SD and pooled from at least two independent experiments (n ≥ 10 cells per experiment). (G–J) Fluorescence images of control (G,I) and SJ1^RQ^KI (H,J) iPSC-derived dopaminergic neuron cultures at days 55–65, expressing GFP-LC3 to label autophagosomes in axons (G,H) and soma (I,J). Scale bars, 10 µm. (K) Quantification of the number of autophagosomes in axons and soma of control and SJ1^RQ^KI dopaminergic neurons, respectively. ****, p < 0.0001 (Mann–Whitney test); n = 3 independent experiments. (L) Electron micrographs of SJ1^RQ^KI neurons showing the presence of large (left two panels) and small (right panels) SV-like organelles within autophagosomes, with representative insets. Scale bars, 500nm.

Given our findings that synaptogyrins, SV2C, and VMAT2 define molecularly distinct dopamine vesicle populations, we next asked whether disruption of synaptic vesicle recycling pathways could preferentially affect these vesicles under PD-associated conditions. To address this question, we examined dopaminergic neurons carrying the PD-associated synaptojanin-1 (SJ1) R219Q mutation (SJ1^RQ^KI; also referred to as R258Q), which impairs clathrin uncoating during synaptic vesicle endocytosis. In mice harboring this mutation, clathrin-coated intermediates accumulate throughout the brain, with dopaminergic terminals in the dorsal striatum exhibiting selective dystrophy of DAT- and tyrosine hydroxylase-positive structures resembling onion-like membrane whirls (Cao et al., 2017; Ng et al., 2023). These dystrophic structures are enriched in SV2C, DAT, VMAT2, and additional synaptic proteins identified in the human PD datasets, including AADC and SNAP25 (Figure 5A), and occasionally contain endosome-like membranes, pointing to profound disruption of vesicle trafficking pathways within dopaminergic terminals(Cao et al., 2017; Lin et al., 2025; Ng et al., 2023).

Interestingly, cultured iPSC-derived dopaminergic neurons carrying the SJ1^RQ^KI mutation did not exhibit overt membrane whirl structures. Instead, these neurons exhibited pronounced clustering of endocytic factors at presynaptic sites (Figure 5D–F), consistent with impaired endocytic turnover, as previously reported in SJ1 mutant neurons ((Cao et al., 2017; Mohd Rafiq et al., 2024). Since the human PD datasets also highlighted alterations in autophagy-related pathways, we next examined whether impaired vesicle recycling in SJ1 mutant neurons was associated with defects in vesicle clearance. Notably, we observed a significant increase in GFP-LC3-positive autophagosomes in both axons and soma of SJ1^RQ^KI dopaminergic neurons at days 55–65 in culture (Figure 5G–K). Electron microscopy revealed that a subset of autophagosomes contained two distinct vesicle populations: small clathrin-coated vesicles (∼40 nm) and larger vesicles ranging from ∼60 to 100 nm in diameter (Figure 5L). It remains unclear whether these small and enlarged vesicles correspond to distinct vesicular transporter populations; future studies will be required to address this question. Together, these observations support the idea that defective endocytic trafficking and vesicle clearance are key features of dopaminergic vulnerability in PD.

## DISCUSSION

Coexpression of synaptophysin or synaptogyrins with synapsin is sufficient to generate clusters of small, SV-like vesicles in reconstitution systems, and loss of these proteins results in enlarged vesicles at synapses(Park et al., 2024). Consistent with this organization, vesicular transporters for glutamate and GABA (VGLUTs and VGAT) preferentially localize to synaptophysin-positive SVs that conform to canonical size constraints of approximately 40 nm (Fujise et al., 2025b; Park et al., 2023).

In contrast, dopamine-containing vesicles loaded by VMAT2 do not follow this organization. Rather than coassembling with synaptophysin, VMAT2 segregates into distinct synapsin-containing condensates that give rise to larger and more heterogeneous vesicle populations(Fujise et al., 2025b). This behavior parallels observations in dopaminergic neurons, where VMAT2- and DAT-positive synaptosomes are relatively depleted of synaptophysin(Asmerian et al., 2025; Zhang et al., 2015) and frequently contain large, irregular vesicles(Lapios et al., 2025; Wildenberg et al., 2021). Moreover, the size of these vesicles resemble those observed in mouse brains lacking all four synaptophysin/synaptogyrin family members(Park et al., 2024), indicating that dopamine vesicles are specified through mechanisms distinct from those governing canonical SV biogenesis.

Given prior evidence that synaptogyrin 3 is present in dopamine vesicles(Egaña et al., 2009; Hobson et al., 2022), we tested whether synaptogyrin recruitment could impose canonical size constraints on VMAT2-positive vesicles. Although synaptogyrins, particularly synaptogyrin 3, associate with VMAT2-positive vesicles and increase vesicle size uniformity, they do not restore canonical SV size (i.e. 40-50 nm). This observation suggests that additional features linked to VMAT2 trafficking contribute to dopamine vesicle sizing. Notably, VMAT2, but not VGLUTs, relies on an AP-3-dependent trafficking pathway, and AP-3 has been shown to recruit lipid-modifying factors such as flippases (e.g., ATP8A1)(Xu et al., 2023). It is therefore possible that vesicles generated through distinct adaptor protein pathways (e.g., AP-3 versus AP-2) differ in lipid composition, which in turn could influence the recruitment or stoichiometry of synaptophysin and synaptogyrin family proteins.

Importantly, our data further identify SV2C as a molecular feature that distinguishes dopamine vesicles from synaptophysin-containing SV populations. Despite the high structural homology among SV2 isoforms, SV2C preferentially associates with VMAT2-positive vesicles in both reconstitution assays and dopaminergic neurons, whereas SV2B favors synaptophysin-defined vesicles when both populations are present. This specialization is consistent with genetic studies showing that loss of SV2C selectively impairs dopamine release and motor behavior (Dunn et al., 2017), in contrast to the essential role of SV2A in general neural viability(Bartholome et al., 2025; Janz et al., 1998). Together, these observations suggest that SV2C is preferentially routed through dopamine-specific trafficking pathways, although the molecular basis of this selectivity remains to be determined.

Finally, our findings in PD–associated SJ1 mutant neurons link dopamine vesicle identity to vesicle clearance pathways. In iPSC-derived dopaminergic neurons carrying the SJ1 mutation, both small and enlarged SV-like vesicles accumulate within autophagosomes, indicating impaired endocytic processing and clearance of multiple vesicle populations. This vulnerability may reflect the atypical size and trafficking properties of dopamine vesicles and provides a potential mechanistic link between vesicle specialization and selective dopaminergic neurodegeneration in PD.

## ACKNOWLEDGEMENTS

N.M.R. acknowledges support from the Deutsche Forschungsgemeinschaft (DFG, German Research Foundation; project number 335549539—GRK 2381) and core funding from the Excellence Strategy at the University of Tübingen. This work was also supported by grants to K.F. from the Japanese Society of Microscopy Young Researchers Research Grant, the Pharmacological Research Foundation, and the Ohsumi Frontier Science Foundation, as well as by grants to A.S. from the Takeda Science Foundation and the Toray Science Foundation. K.F. and A.S. acknowledge support from the Center for Biomedical Research and Education, Kanazawa University. Additional support was provided in part by grants to Pietro De Camilli (Yale University, USA), for which we are grateful. We thank Peng Xu (Yale University, USA) for assistance with mouse husbandry.

## AUTHOR CONTRIBUTIONS

K.F. and N.M.R. conceptualized the project. K.F. and N.M.R. designed and performed most experiments. N.K. performed the AlphaFold analyses, while J.M., H.K., T.T., and A.S. provided experimental assistance. N.M.R. and K.F. wrote the manuscript, and N.M.R. coordinated the project. All authors read and approved the final manuscript.

## METHOD

### Plasmids and constructs

All SV2 isoform constructs used in this study were based on human codon-optimized sequences and cloned using the Gateway recombination cloning system. The following donor constructs were obtained from the RESOLUTE consortium SLC collection on Addgene: pDONR221_SV2A (#132257), pDONR221_SV2B (#132281), and pDONR221_SV2C (#132297). These constructs were C-terminally tagged with EGFP by recombination into the destination vector pDEST-eGFP-N1 (#31796).

The following plasmids were previously generated in the laboratory of Pietro De Camilli: untagged synaptophysin, synaptogyrin 1-myc, synaptogyrin 2-myc, synaptogyrin 3-myc, untagged synaptogyrin 3, untagged synapsin 1a, mCherry-synapsin 1a, KDEL-HRP and EGFP-LC3. VMAT2-GFP, VMAT2-FLAG, VGAT-FLAG, and VGLUT2-GFP constructs were obtained from the laboratory of Nisha Mohd Rafiq; sequence information for these constructs has been described previously (Fujise et al., 2025b).

### Cell culture and DNA transfection

COS7 cells were grown in DMEM (Thermo Fisher Scientific) supplemented with 10% FBS (Thermo Fisher Scientific) and 1% penicillin-streptomycin. Cells were kept at 37 °C with 5% CO2 in an enclosed incubator. For transfection of COS7 cells, 1 μl Lipofectamine™ 2000 Transfection Reagent (Invitrogen) was used with the respective plasmids and visualized within 24–48 h.

The following iPSC lines were obtained from the iNDI consortium and genome-edited by Jackson Laboratories (JAX): KOLF2.1, KOLF2.1 (with the NGN2 cassette at the AAVS locus; RRID:CVCL_D1KS), used for the i3Neurons experiments) and KOLF2.1 SJ1^RQ^KI (R219Q): clones A09 and B02. For the maintenance of iPSCs in culture, iPSCs were cultured on Geltrex (Life Technologies) coated dishes and maintained in Essential 8 Flex media (Thermo Fisher Scientific). The Rho-kinase (ROCK) inhibitor Y-27632 (EMD Millipore, 10 μM) was added to Essential 8 Flex media on the first day of plating and replaced with fresh media without ROCK inhibitor on the following day.

For i^3^neuronal differentiation, iPSCs were differentiated into cortical-like i^3^Neurons according to a previously described protocol based on the doxycycline inducible expression of Ngn2(Fernandopulle et al., 2018). A detailed protocol can be found at https://www.protocols.io/view/culturing-i3neurons-basic-protocol-6-n92ld3kbng5b/v1. For the differentiation of iPSCs to DA neurons, we used the following protocols described in (Kriks et al., 2011) and (Bressan et al., 2021). A detailed protocol can be found at https://doi.org/10.17504/protocols.io.dm6gp39m8vzp/v1.

For both i^3^Neuron and DA neuron transfections, plasmids were transfected with 4 μl of Lipofectamine™ Stem Transfection Reagent (Invitrogen) and visualized at least 48 h later.

### Immunofluorescence, live imaging and fluorescent microscopy

For all imaging, cells were seeded on glass-bottom matTek dishes (MatTek corporation) or ibidi dishes (ibidi GmbH). For immunofluorescence visualization, cells were fixed with 4% (v/v) paraformaldehyde (Electron Microscopy Sciences) in 1x phosphate-buffered saline (PBS) for 20 min followed by three washes in PBS. Cells were permeabilized with 0.25–0.5% (v/v) Triton X-100 in PBS for 5 min followed by three washes in PBS. Cells were further blocked for 30 min in 5% bovine serum albumin (BSA, Sigma-Aldrich) in PBS and then incubated overnight at 4 °C with the primary antibodies listed in Supplementary Table 1. Subsequently, cells were washed with PBS thrice the following day and incubated with Alexa Fluor-conjugated secondary antibodies (Thermo Fisher Scientific) for 1 h at room temperature, followed by three washes in PBS.

Fluorescence imaging was performed using spinning-disk and laser-scanning confocal microscopy systems. Imaging of COS7 cells was performed using either an inverted spinning-disk confocal microscope (Dragonfly 200; Oxford Instruments) equipped with a Zyla 5.5 cMOS camera and controlled by Fusion software, or the LSM980 with Airyscan 2. Excitation wavelengths ranging from 405 to 640 nm were used as appropriate. Images were acquired using a Plan Apo 60× oil-immersion objective (NA 1.45).

For autophagosome imaging in neurons, cells were imaged live using either a spinning-disk confocal microscope (CSU-W1 SoRa; Nikon) controlled by NIS-Elements software, or a laser-scanning confocal microscope (LSM980; Zeiss) equipped with Airyscan 2 detection.

### EM sample preparation and CLEM

For EM imaging, COS7 cells were plated on glass-bottom MatTek dish (P35G-1.5-14-CGRD) and transfected as described above. Cells were fixed in 4% PFA then washed with PBS before fluorescence light microscopy imaging. Regions of interest were selected and their coordinates on the dish were identified using phase contrast. Cells were further fixed with 2.5% glutaraldehyde in 0.1 M sodium cacodylate buffer, postfixed in 2% OsO_4_ and 1.5% K_4_Fe(CN)_6_ (Sigma-Aldrich) in 0.1 M sodium cacodylate buffer, en bloc stained with 2% aqueous uranyl acetate, dehydrated in graded series of ethanols (50%, 75%, and 100%), and embedded in Embed 812. Cells of interest were relocated based on the pre-recorded coordinates. Ultrathin sections (50–60 nm) were post-stained with uranyl acetate substitute (UranyLess, EMS), followed by a lead citrate solution. Sections were observed in a Talos L 120 C TEM microscope at 80 kV, images were taken with Velox software and a 4k × 4 K Ceta CMOS Camera (Thermo Fisher Scientific).

### Horseradish Peroxidase (HRP) assay

For HRP uptake assay, COS7 cells were incubated with 15 μg/ml HRP conjugated CTX (Thermo Fischer Scientific) at 37 °C for 1h. To stain ER, COS7 cells were transfected ER-HRP. HRP reaction was carried out with diaminobenzidene (Sigma-Aldrich, D-5637) (0.5 mg/ml) and H_2_O_2_ (JTBaker, 2186-01) (0.01%) in 0.1 M ammonium phosphate buffer (pH7.4) after the glutaraldehyde fixation step. Ultrathin sections (50–60 nm) were observed in Talos L120C TEM microscope at 80 kV. Images were taken with Velox software and a 4k × 4 K Ceta CMOS Camera (Thermo Fischer Scientific). Except when noted, all EM reagents are from Electron Microscope Sciences.

### Transcriptomic analysis of postmortem PD striatum

Bulk RNA-seq data from postmortem caudate and putamen of 35 PD patients and 40 neurologically normal controls were obtained from the Gene Expression Omnibus (accession GSE205450). Differential gene expression results were used as provided from Irmady et al. (2023). Genes were considered significantly differentially expressed at an adjusted p-value < 0.05 (Benjamini-Hochberg FDR correction). Volcano plots and heatmaps were generated in R using ggplot2. All visualizations were generated in R (version 4.x) using ggplot2, ggrepel, SynGO (Koopmans et al., 2019), and patchwork.

### Statistical analysis

The methods for statistical analysis and sizes of the samples (n) are specified in the results section or figure legends for all quantitative data. Comparisons between two groups were performed using Student’s t-test, Kruskal-Wallis test or the Mann–Whitney test, whereas comparisons among three groups were performed using one-way ANOVA. Differences were accepted as significant for P < 0.05. Prism version 10 (GraphPad Software) was used to plot, analyze and represent the data.

### Data availability

All data generated or analyzed during this study are included in this published article (and its Supplementary Material). Raw datasets generated during and/or analyzed during the current study are available from the corresponding author on request.

---

**Supplementary Figure 1.**
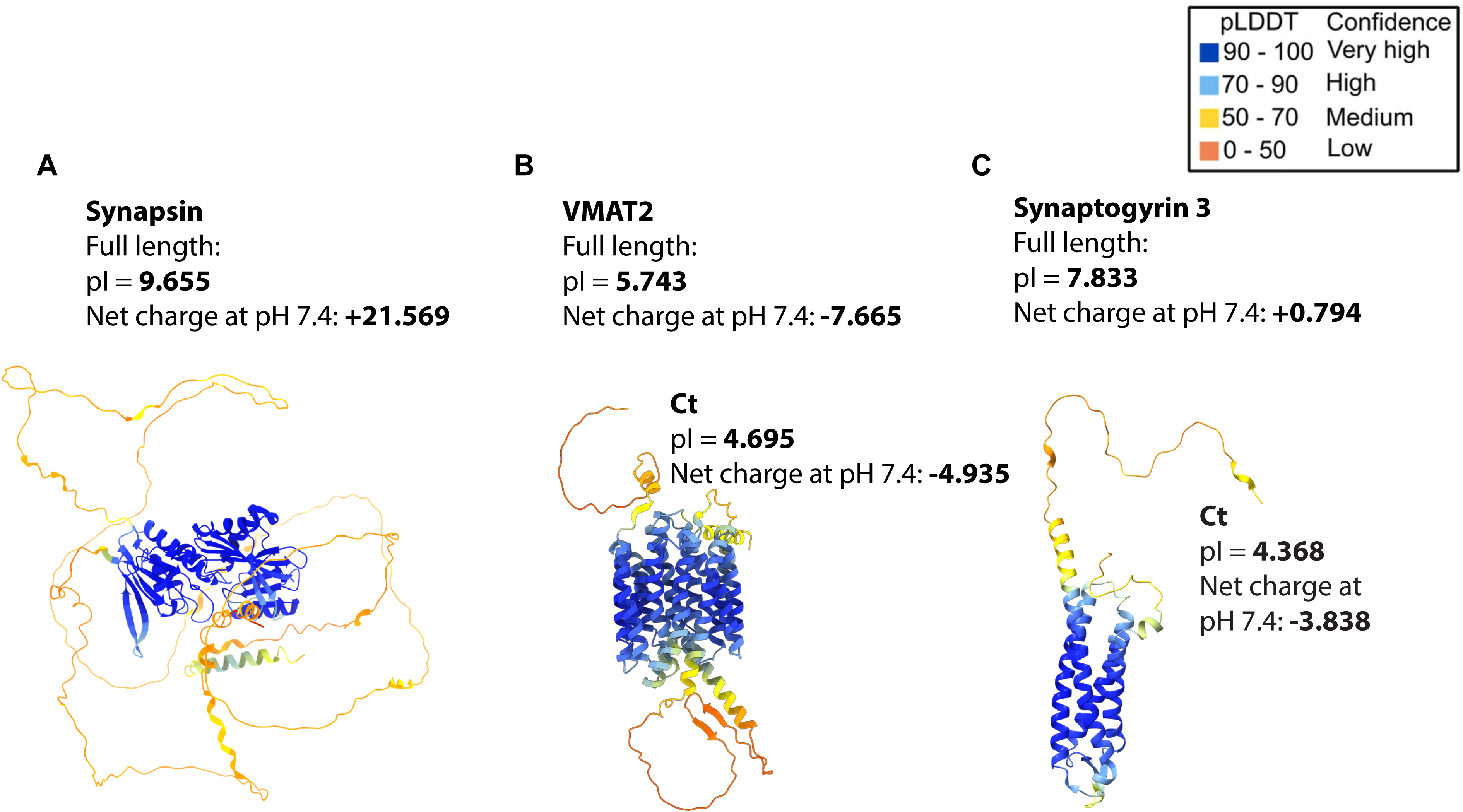
Predicted structures and electrostatic properties of synapsin, VMAT2, and synaptogyrin 3. (A–C) AlphaFold-predicted structures of full-length human synapsin (A), VMAT2 (B), and synaptogyrin 3 (C), together with their isoelectric point (pI) values and calculated net charges for the full-length proteins and their C-terminal regions. AlphaFold DB accession numbers are synapsin (AF-P17600-F1), VMAT2 (AF-Q05940-F1), and synaptogyrin 3 (AF-O43761-F1). Structures are colored according to pLDDT (predicted Local Distance Difference Test) scores, indicating per-residue prediction confidence (top right). Theoretical isoelectric points were calculated using the ProtPi Protein Tool (https://www.protpi.ch/Calculator/ProteinTool). Note that both VMAT2 and synaptogyrin 3 possess negatively charged C-terminal regions that could potentially interact with the highly basic C-terminal domain of synapsin.

**Supplementary Figure 2.**
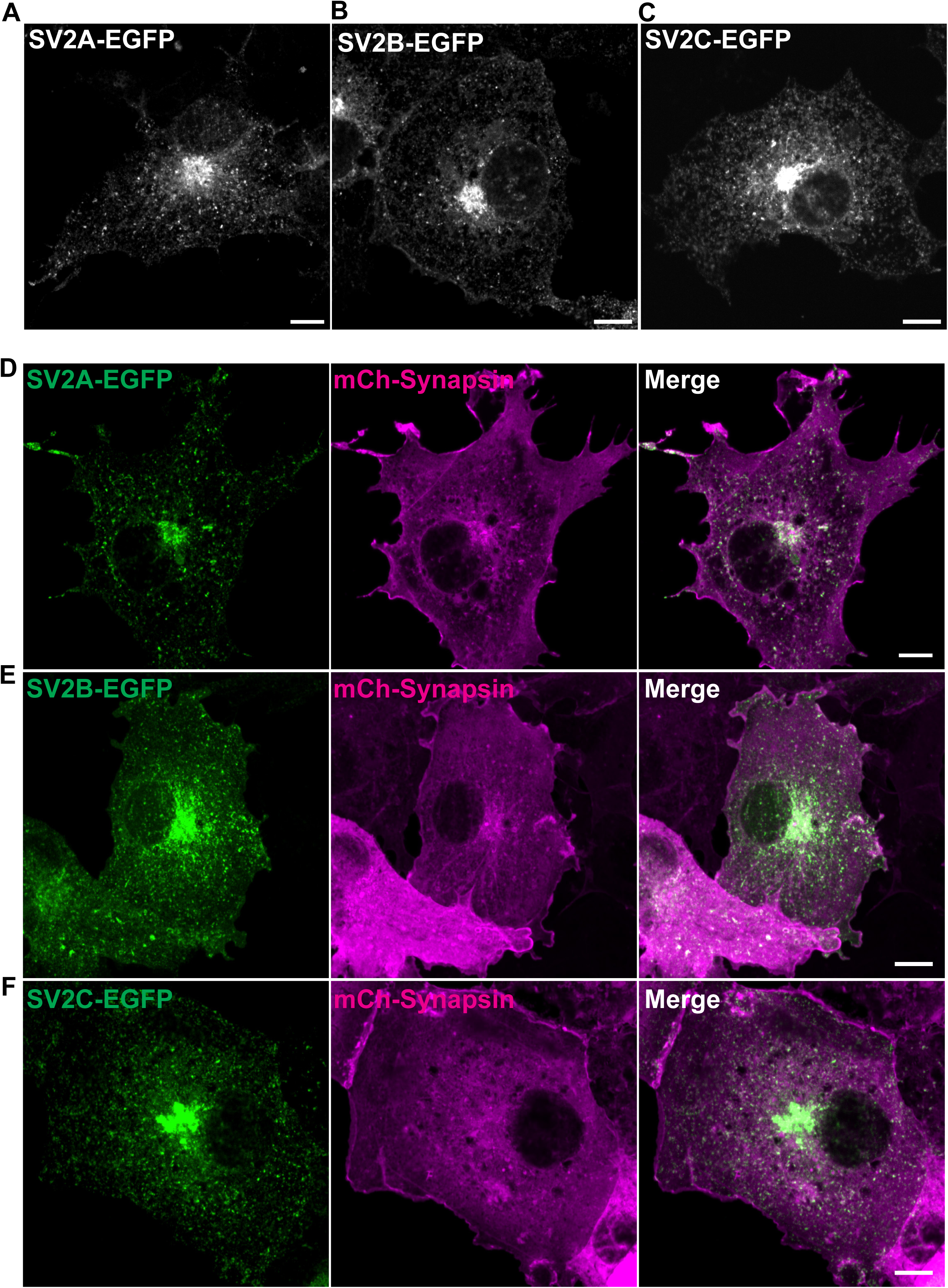
SV2 isoforms do not induce synapsin-dependent condensates in COS7 cells. (A–C) Representative fluorescence images of COS7 cells transfected with SV2A-GFP (green; A), SV2B-GFP (green; B), or SV2C-GFP (green; C), showing Golgi-associated and vesicular punctate staining. (D–F) COS7 cells co-transfected with mCherry-synapsin (magenta) together with SV2A-GFP (green; D), SV2B-GFP (green; E), or SV2C-GFP (green; F) do not form condensate-like structures. Scale bars, 10 µm.

**Supplementary Figure 3.**
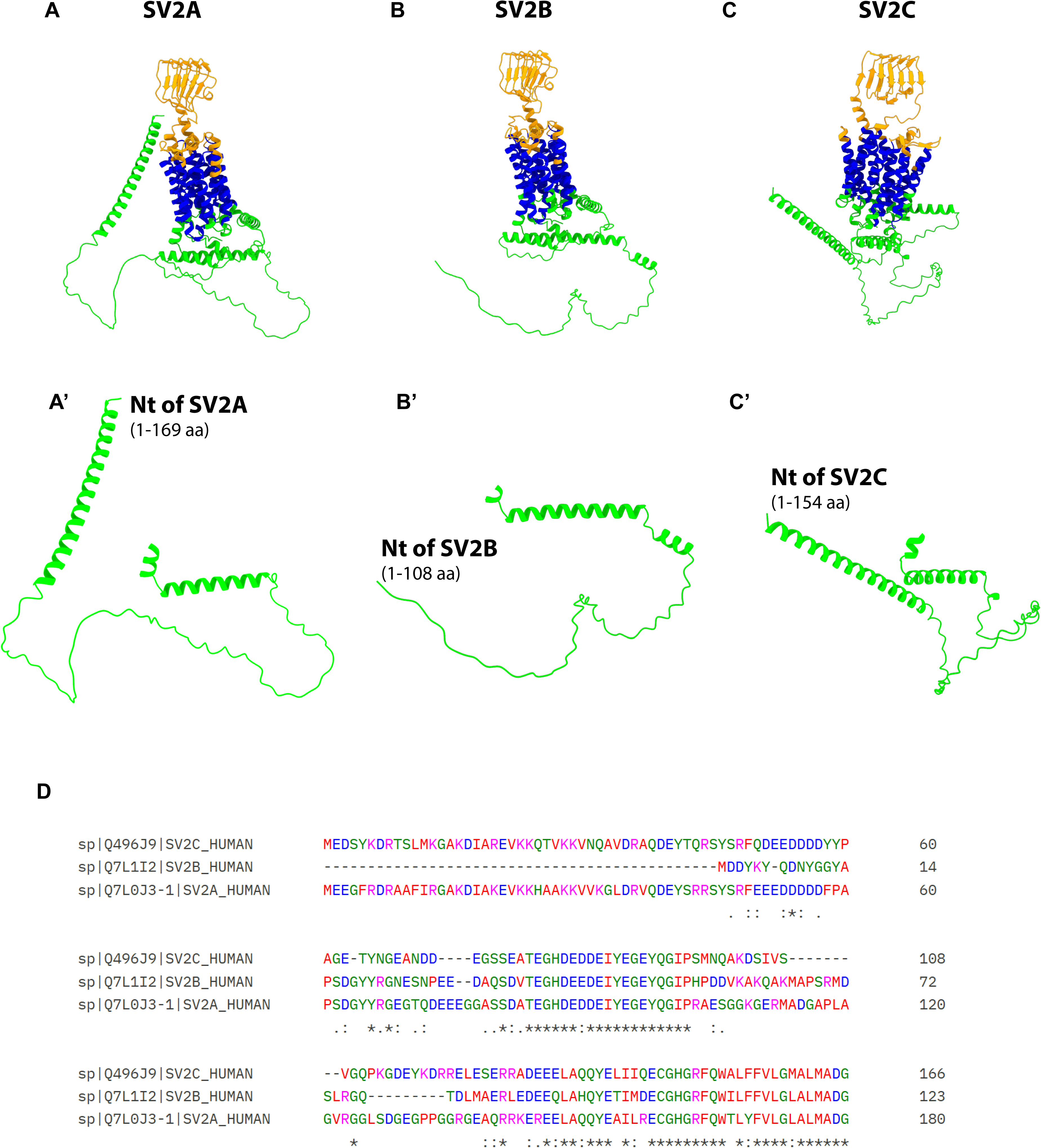
Predicted structural features and N-terminal divergence of SV2 isoforms. (A–C) AlphaFold2-predicted structural models of full-length human SV2A (AF-Q7L0J3-F1, A), SV2B (AF-Q7L1I2-F1, B), and SV2C (AF-Q496J9-F1, C), obtained from UniProt. Protein topology annotations were retrieved from UniProt and used to color structural domains, with transmembrane helices shown in blue, luminal regions in orange, and cytoplasmic regions in green.(A’-C’) Isolated N-terminal cytoplasmic domains of SV2A (A’), SV2B (B’), and SV2C (C’), respectively, generated by masking non–N-terminal regions based on UniProt-defined boundaries; the length of each region is indicated in amino acids (aa). (D) Multiple sequence alignment of the N-terminal cytoplasmic regions of SV2 isoforms generated using Clustal Omega. Note the greater similarity between the N-termini of SV2A and SV2C compared with SV2B.

**Table S1.** List of antibodies/dyes used in this study.

| **Target Protein** | **Company;**<br>**Catalog**<br>**number** | **Antibody**<br>**species** | **Working dilution for**<br>**immunofluorescence** | **Working**<br>**dilution for**<br>**immunoblotting** |
| --- | --- | --- | --- | --- |
| Dopamine transporter | EMD Millipore;<br>MAB369 | Rat | 1:500 | N/A |
| SV2C | Synaptic<br>Systems;<br>119202 | Rabbit | 1:500 | N/A |
| Synapsin I | Synaptic<br>Systems;10611 | Mouse | 1:250 | N/A |
| Synapsin I | Synaptic<br>Systems;<br>106103 | Rabbit | 1:250 | N/A |
| Synaptophysin | Synaptic<br>Systems;<br>101002 | Rabbit | 1:500 | N/A |
| Synaptophysin | Synaptic<br>Systems;<br>101001 | Mouse | 1:250 | N/A |
| FLAG | Sigma-Aldrich;<br>F1804 | Mouse | 1:1000 | N/A |
| Myc | Sigma-Aldrich;<br>9E10 | Mouse | 1:1000 | N/A |
| Amphiphysin-2 | Sigma-Aldrich;<br>99D | Mouse | 1:250 | N/A |

